# The VersaLive platform enables pipette-compatible microfluidic mammalian cell culture for versatile applications

**DOI:** 10.1101/2021.07.02.450869

**Authors:** G. M. Nocera, G. Viscido, S. Brillante, S. Carrella, D. di Bernardo

**Affiliations:** Telethon Institute of Genetics and Medicine (TIGEM), Via Campi Flegrei 34, 80078 Pozzuoli (NA), Italy; Department of Electrical Engineering and Information Technology, University of Naples Federico II, Naples, Italy; Department of Chemical, Materials and Industrial Production Engineering, University of Naples Federico II, Naples, Italy

## Abstract

Microfluidic-based cell culture allows for precise spatio-temporal regulation of microenvironment, live cell imaging and better recapitulation of physiological conditions, while minimizing reagents consumption. Despite their usefulness, most microfluidic systems are designed with one specific application in mind and require specialized equipment and expertise for their operation. All these requirements prevent microfluidic-based cell culture to be widely adopted. Here, we designed and implemented a versatile and easy-to-use perfusion cell culture microfluidic platform for multiple application (VersaLive) requiring only standard pipettes. Here, we showcase the multiple uses of VersaLive (e.g., time-lapse live cell imaging, immunostaining, cell recovery, cell lysis) on mammalian cell lines and primary cells. VersaLive can replace standard cell culture formats in several applications, thus decreasing costs and increasing reproducibility across laboratories. The layout, documentation and protocols are open-source and available online at https://versalive.tigem.it/.

## Main

Micro-scaled systems can be designed to accommodate experimental requirements that are difficult to meet in standard mammalian cell culture or utterly impossible. In general, the use of microfluidics in cell culture allows for precise control of the extrinsic factors (e.g., nutrients, drug treatment, sample confinement) while better mimicking physiological conditions and, thanks to the reduced volumes, minimizing reagents consumption [Kwon *et al.*, 2017; Gagliano *et al.*, 2019; Sohn *et al.*, 2020]. Some of the recent applications of microfluidics in mammalian cells are high throughput single-cell sequencing [Klein *et al.*, 2015; Macosko *et al.*, 2015], parallelized drug screening [Schuster *et al.*, 2020], temporal modulation of treatments [Kolnik, Tsimring and Hasty, 2012], and automated feedback control of biological processes [Postiglione *et al.*, 2018]. However, microfluidic approaches are usually designed for specific applications and often involve complex multi-layer microfabrication processes, specialized equipment and extensive expertise that currently greatly limit their adoption by academic and industrial research laboratories.

Here, we designed and developed a versatile microfluidic device, which we named VersaLive, to enable mammalian cell culture, imaging and cell manipulation. To enable wide adoption of VersaLive and to make it easy to transfer standard experimental protocols to microfluidics, we designed VersaLive so that all the operations can be carried out by manual pipetting. We demonstrated culture of cell lines and primary cells and application of a variety of protocols from immunostaining to cell lyses or retrieval.

The device is shown in Figure 1; it consists of a gas-permeable elastomer (polydimethylsiloxane, PDMS) patterned with channels that develop on one single layer bound onto a microscopy glass coverslip. The device is engineered over one single layer to allow fast and easy replication of the technology also in laboratories without previous expertise in microfluidics. As shown in Figure 1a, the platform consists of five 250 μm-wide chambers. Each chamber is connected on one side to a main common channel and on the other side to an independent input channel. The input channel is doubled to increase robustness against clogging. Channels are accessed through ports (Figure 1a) that act as reservoirs where inputs (e.g. growth medium, drugs, reagents) can be changed. Hydrostatic pressure alone drives the flow from the ports to the chamber, thus doing away with pumps, motors or pressure regulators. Cell filters are embedded at one side of each culture chamber to contain cells within the chamber by preventing anything larger than 5 μm in width to go through the filter. A fluidic resistor between the culture chamber and the input channels decreases the flow velocity of the cells during loading and prevents shear stress onto the cells during perfusion. The flow deflectors help in spreading the chemical input over the whole chamber area for a uniform and consistent delivery.

**Figure 1:**
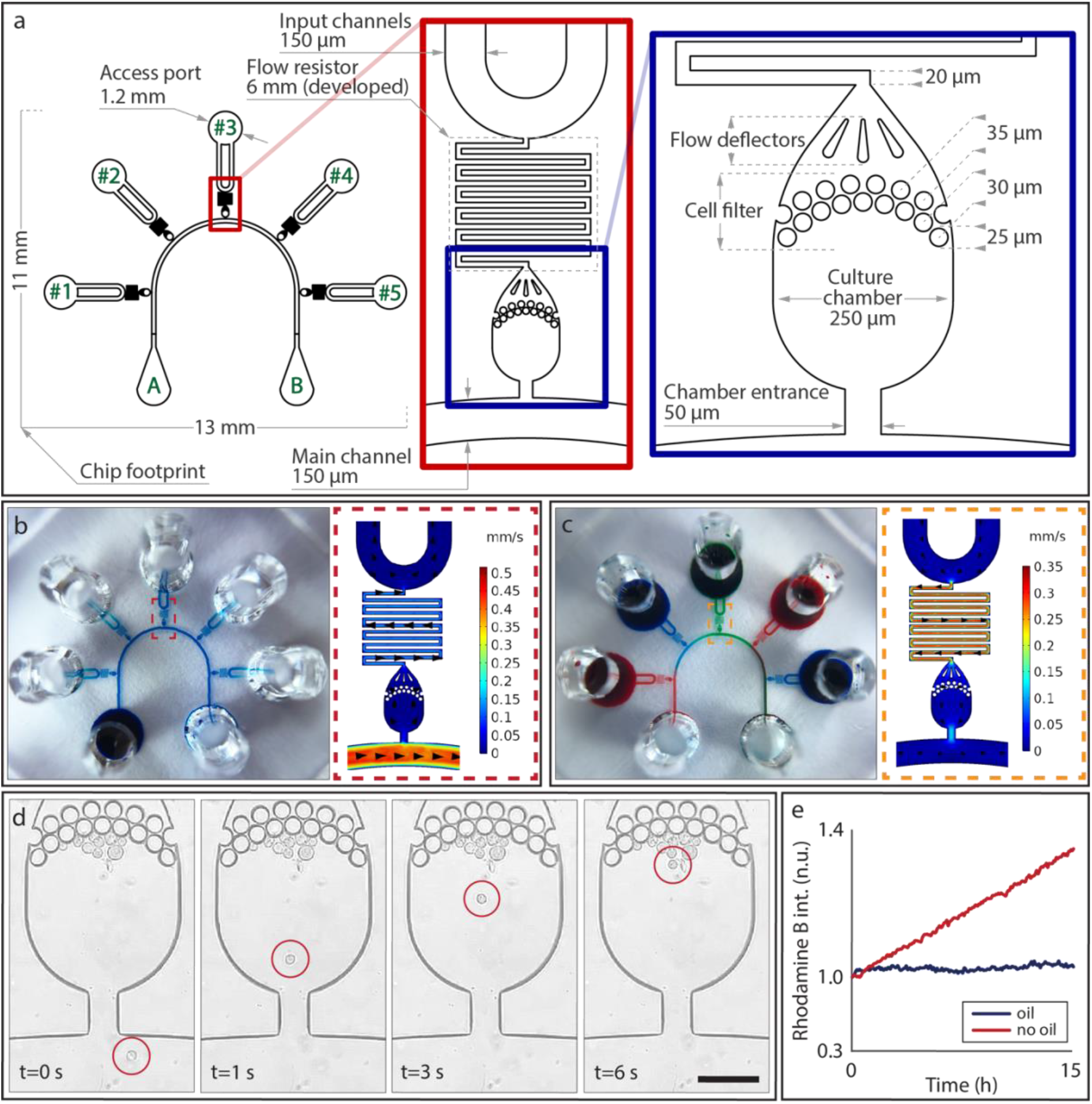
Outline of the VersaLive microfluidic platform. **(a)** Layout of the microfluidics device with five independent chambers. On one side, all chambers are connected to the main channel that develops from port A to port B of the schematics. Fluids can be fed to each chamber independently by a dedicated port (ports #1 to #5 on the schematics). The flow resistor serpentine prevents shear stress to the cells during the utilization of the device. The flow deflectors ensure an even distribution of the fluid across the whole area of the chamber. **(b, c)** Modes of operation of VersaLive and their finite element simulations in COMSOL Multiphysics software (dashed inset). In the “perfusion” single-input mode (b) the main channel is exploited to deliver the same medium to all chambers at the same time. Multi-input mode (c) exploits the dedicated ports to deliver different inputs (i.e. media, chemicals) to each chamber avoiding cross-contaminations. **(d)** During cell loading, the cell suspension flows through the main channel and into the chambers. Once in the chamber, cells slow down and are eventually stopped by the filter features as predicted in the velocity profile of the simulation. Scale bar 150 μm. **(e)** Solvent evaporation at the reservoirs was observed to increase the solute concentration of the channel content during perfusion. The effect was completely removed by the addition of a layer of mineral oil at the reservoirs.

All operations in VersaLive are carried out by manually pipetting content directly in and out of the ports that effectively act as reservoirs. VersaLive can be operated in either of two ways: *single-input mode*, where the same input (e.g., cell growth medium) is delivered to all cell chambers (Figure 1b) or the *multi-input mode*, where a different input can be delivered to each of the 5 chambers (Figure 1c). Single-input mode is set up by pipetting a volume of up to 20 μL in the reservoir connected to the main channel (port A in Figure 1b) while leaving all other reservoirs empty. This configuration was called the “*perfusion” single-input mode*, as one constant flow from the filled reservoir to the cell chambers will be present. A variant of the single-input mode is obtained if all reservoirs are filled with the same volume. In this case, no flow will be present in the device, giving rise to a static cell culture. This *“static” single-input mode* of operation is particularly convenient during the cell adhesion phase, when the cells are not fully attached to the chip and a flow would risk displacing them.

In the case of the multi-input mode, all input reservoirs are filled while the main channel ports are left empty (Figure 1c). This configuration creates an active flow across each chamber that it is strong enough to prevent backflow from the main channel, but sufficiently slow to prevent shear stress to the cells. We verified that a volume of 20 μL will suffice for about 24 hours of constant perfusion. Multi-input mode is applicable, for instance, when different concentrations of a small molecule or antibody need to be tested in parallel, as in drug screening or immunostaining. For each configuration, the hydrostatic pressure difference across the chip results in the velocity profile shown by the finite element simulation in the dashed insets (Figure 1b, c).

Independently of the chosen mode of operation, the chip is always initialized by loading cells into the chambers using the “perfusion” single-input mode, as shown in Figure 1d. Time-lapse acquisition of the cell loading step were used to estimate a cell loading velocity of 150 μm per second at the entrance of the chamber and then it takes an additional 5 seconds to further travel 150 μm before eventually stop against the filter, in line with the velocity profile obtained in simulations and shown in Figure 1b, c.

Exposed to the external environment, the small volume of the reservoirs tends to evaporate even when kept in a cell incubator, as shown in Figure 1e. This effect, however, can be completely eliminated by pipetting 2.5 μL of mineral oil to the reservoirs, as shown in Figure 1e where an increase over time of the solute concentration in the reservoir was observed in the absence of oil, but completely abolished in its presence.

The ability of the microfluidic platform to deliver independent chemical inputs to each of the five culture chambers was validated by using a CHOP::GFP tagged CHO-K1 cell line [Crespillo-Casado *et al.*, 2017]. CHOP is a transcription factor whose expression is induced by the activation of the Integrated Stress Response (ISR) pathway and upon endoplasmic reticulum (ER) stress. Cells were loaded into the chambers and kept overnight in incubator with the static single-input mode in standard growth medium to ensure cell adhesion to the glass surface. The following day, all the reservoirs were emptied by pipetting, and the chip was switched to the multi-input mode of operation by adding fresh medium to the reservoirs connected to chambers #2 and #4, while tunicamycin dissolved in fresh medium (0.5 μg/mL) was added to chamber #1, #3 and #5, as shown in Figure 2a. Tunicamycin is a potent inducer of ER stress by blocking N-glycosylation of proteins and thus should induce CHOP::GFP expression in the reporter cells. The VersaLive chip was then placed on an inverted fluorescence microscope and each chamber was imaged for 20 hours at 15 minutes intervals. As shown in Figure 2a–c, at the end of the treatment, only cells exposed to tunicamycin expressed CHOP::GFP, confirming no cross-flow among chambers. Moreover, as an additional check for the proper function of the device, we added rhodamine B to the input reservoir of chamber #3. This red fluorescent dye has no toxic effects on cells. As expected, red fluorescence was detected only at the outlet of chamber #3 but was absent in the other four chambers. The expression of the CHOP::GFP in time, reported in Figure 2b, shows appreciably higher expression in the treated samples already after 8 hours of exposure to tunicamycin. To quantify the difference in ER stress among the chambers, the last time point of the treatment was analyzed with single cell resolution. The 20-hour tunicamycin exposure resulted in a 15-fold increase in the CHOP::GFP expression (Figure 2c).

**Figure 2:**
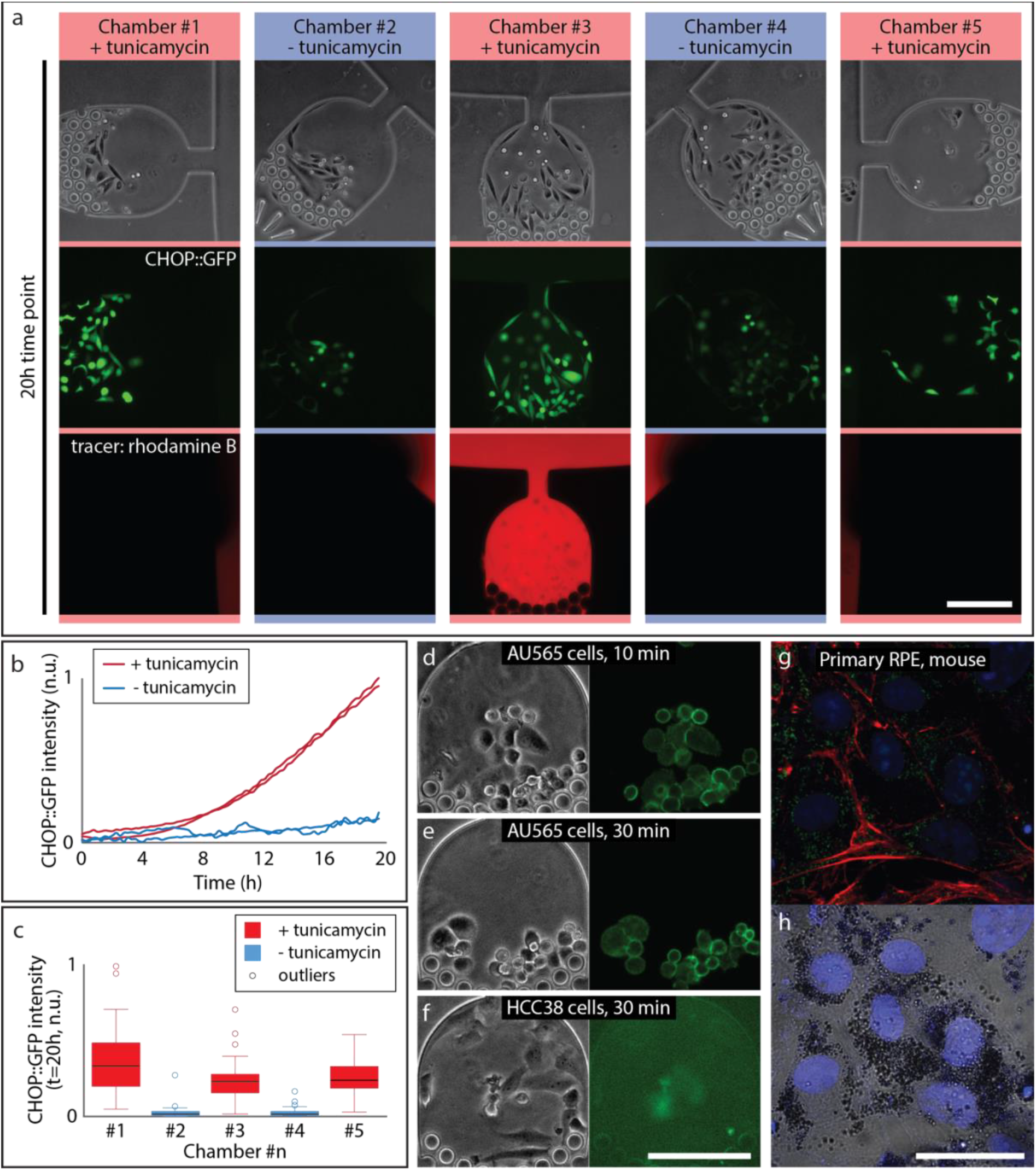
Applications of VersaLive. **(a)** CHO-K1 cells in the indicated chambers were exposed to tunicamycin treatment (0.5 μg/mL for 20 hours) and stress response was measured by the intensity of CHOP::GFP fluorescence. The third row shows rhodamine B that was used as flow tracer to check proper flow in the platform throughout the time-lapse acquisition. Scale bar 150 μm. **(b)** Time-course of the average CHO::GFP fluorescence across cells in chambers #1, #2, #3, #4 (15 minutes time resolution). **(c)** Bar-plot of the quantification of the CHO::GFP fluorescence in single cells following 20 hour of tunicamycin treatment in the indicated chambers. **(d-f)** On-chip immunofluorescence of HER2 membrane receptor. Immunostaining following either 10 minutes perfusion or 30 minutes of perfusion of AU565 cells (d, e) and HCC38 cells (f). Scale bar 150 μm. **(g, h)** Culture of primary mouse RPE cells. On-chip chemical fixation and immunostaining, labeling of citrate synthase (green), f-actin (red) and cell nuclei (blue). The retina cells were able to adhere to the chip while retaining their characteristic dark pigment. Scale bar 30 μ.m

The use of VersaLive is not exclusive to live cell imaging. The single- and multi-input modes of operation can be exploited to implement any protocol of interest such as delivering fixing agents (e.g., paraformaldehyde) and labelled antibodies directly to the cells. Washing and staining steps of established immunostaining protocols can be replicated with the advantage of an effective reagent consumption in the nanoliter range and with operation times cut down from hours to minutes [Kwon *et al.*, 2017]. The human epidermal growth factor receptor 2 (HER2) is a membrane protein among the biomarkers targeted in immunohistochemistry for the diagnosis of breast cancer. Cancer cells expressing HER2 (HER2+) are more proliferative but result in a better prognosis while those not expressing HER2 (HER2−) are more resistant to cytotoxic chemotherapy [Jordan *et al.*, 2016]. For this reason, discrimination between these two kinds of tumors is important to decide the therapeutic strategy that will be carried out. Here, HER2+ (AU565) and HER2-(HCC38) breast cancer cell lines were used to validate immunofluorescence protocol in VersaLive. The two types of cells were cultivated on two different VersaLive chips, chemically fixed and fluorescently labeled via anti-HER2 immunostaining directly on the device. As shown in Figure 2d–f, immunostaining in microfluidics was efficient and resulted in the correct membrane localization of the HER2 receptor. Immunostaining was already visible following 10 minutes perfusion of the cell chambers with the fluorescent antibody and results were indistinguishable from samples stained for 30 minutes with the same method.

VersaLive also allows direct access to the cells by complete removal of the PDMS from the glass slide where it was reversibly bonded (Supplementary Figure 1a, b). We envision this feature to be useful for cell re-plating, cell sequencing and all applications that require full recovery of the sample. To demonstrate that the PDMS removal operation does not damage the cells, in Supplementary Figure 1c, d, the same field of view was retrieved and compared between the live imaged and the chemically fixed sample.

Retrieval of rare cells (less than 1000 cells per milliliter of sample) are of great importance for the diagnosis of diseases (e.g., circulating tumor cells) or for their recovery from intrinsically minute samples (e.g., small biopsies, small animal organs). Over the years, multiple microfluidic systems have been developed to isolate and concentrate specific types of cells from a bulk sample according, for example, to their phenotype [Chen *et al.*, 2014]. With VersaLive, it is possible to handle a few cells per chamber by simply pipetting microliters of cell suspension and cell medium in and out of the access ports. This characteristic makes VersaLive ideal to grow and experiment with rare cells. Due to the minute size of mouse retina, primary mouse retina pigment epithelium (RPE) cells were used here as rare cell model to demonstrate the target application of VersaLive. RPE cells retrieved from fresh tissue were able to adhere to the glass surface of the chip (Figure 2g, h). Cells were then fixed and immunostained in chip and imaged (Figure 2h).

If required, VersaLive can be also used to recover the chamber content in a selective and precise fashion. To test this possibility, we grew HeLa cells overnight and then the medium in each chamber was sequentially replaced with cell lysis buffer starting with cell chamber #1. As shown in Supplementary Figure 2, when lysing cells in chamber #1, the content of chamber #2 is not affected by the lysis process, and so on for all the chambers. This ability of VersaLive can be used to retrieve DNA and RNA content at specific times during a microscopy time-lapse experiment.

Additional protocols can be easily adapted to VersaLive by rescaling concentrations used macroscopic culture system, such as DNA transfection, with the added advantage of reducing costs and speeding up reactions.

VersaLive offers an easy-to-adopt microfluidic platform for mammalian cells, it enables the choice between perfusion and static cell culture while not requiring any external components to function. VersaLive also allows direct access to the cultured cells with the benefit of reducing the consumption of reagents and plasticware.

The VersaLive technology is open-source and all protocols and materials are available online (https://versalive.tigem.it/) and on protocols.io.

## Methods

### Design and computer simulations of the device

The VersaLive microfluidic platform was designed using Autodesk Fusion 360. The velocity profile of the chip was modelled using the finite element method (COMSOL Multiphysics 5.4). In the simulated utilization modes (**Error! Reference source not found.**), 0.5 mbar of pressure was applied at every inlet while the outlets were set at zero pressure. Flow direction in the simulation plots was indicated using arrows whose dimensions reflect the magnitude of the velocity in a logarithmic scale. A 2D CAD model of the chip was exported into Autodesk AutoCAD 2020 to design the printable photolithography mask.

### Fabrication of the VersaLive microfluidic platform

Microfluidic chips were fabricated by using a combination of standard photolithography and soft lithography procedures [Ferry, Razinkov and Hasty, 2011]. The master mold was microfabricated via mask photolithography of SU-8 negative photoresist (SU-8 3035, Kayaku Advanced Materials Inc.) on silicon wafer substrate. The photoresist was spin coated to reach a final thickness of 25 μm and processed following the guidelines of the manufacturer. Feature height was confirmed by measurement via optical microscopy of the cross-section of a silicone replica of the channels. Before the first utilization of the master, the passivation of its surface is required to facilitate the release step during soft lithography. The master was passivated by vapor deposition of perfluorosilane (1H,1H,2H,2H-perfluorooctyl-trichlorosilane, Merck kGaA). Specifically, the master was placed in a desiccator with a small vial containing 20 μL of perfluorosilane. Vacuum was then applied overnight to allow the perfluorosilane to evaporate and to react with the surface of the silicon wafer, forming a covalently bound super-hydrophobic coating.

The passivated master was then used as a mold for the soft lithography part of the fabrication process of the microfluidic device. The VersaLive microfluidic platform is formed by a silicone elastomer (polydimethysiloxane, PDMS) bonded to a glass slide. The elastomer base of the PDMS was thoroughly mixed with the curing agent in a 10:1 ratio as reported by the datasheet of the manufacturer (Sylgard 184, Dow Corning). Air bubbles were removed from the uncured polymer applying vacuum to the mix for 2 hours or until no bubbles were visible. The mix was then poured onto the master mold for a final thickness of about 5 mm. If required, vacuum was applied a second time to remove the bubbles formed during the pouring step. The mold with the uncured PDMS were then placed in oven at 80°C for a minimum of 2 hours to accelerate the crosslink of the polymer mix. Once cured, the PDMS was peeled off the master mold. The PDMS slab was then placed with the pattern features facing up to cut out the single chips. Similarly, access ports were opened using a 3-mm biopsy punch (ref. 504649, World Precision Instruments).

The channels of the PDMS chips were sealed by plasma bonding the chips to round glass cover slide (30 mm in diameter, thickness no. 1, Marienfeld). Glass slides and PDMS chips were first cleaned from dust particles using adhesive tape. For the surface activation of glass slides and PDMS chips, 85W air plasma at 0.4 mbar or lower (ZEPTO version B, Diener electronic GmbH & Co. KG) were used. For the permanent or reversible bonds, 30 or 10 seconds of air plasma were used, respectively.

The chip was then placed in an aluminum lens tube (SM30L05, ThorLabs) used as chip holder. The lens tube allowed the safe transfer of the chip during the experimental workflow (e.g. workbench, incubator, microscope). To allow an easy and reliable image acquisition procedure, the Nikon microscope was equipped with a custom holder for the lens tubes made of 2-mm laser-cut acrylic.

### Cell culture

HeLa WT, together with AU565 and HCC38 breast cancer cell lines were purchased from ATCC and grown in RPMI 1640 (without L-glutamine, EuroClone) cell medium. CHO-K1 cells [Crespillo-Casado *et al.*, 2017] were grown in F-12 (Gibco) cell medium. Both cell media were supplemented with 10% fetal bovine serum (FBS, EuroClone), 1% L-glutamine (EuroClone) and 1% penicillin-streptomycin (EuroClone). Cells were maintained in a cell incubator at 37°C, 100% humidity and 5% CO_2_ atmosphere. Prior to the loading of a microfluidic device, the content of a T25 flask at 90% confluence was collected according to the following procedure. First, the cell medium in use was removed and the cells were rinsed with 2 mL of phosphate-buffered saline (PBS, EuroClone). To detach the cells from the flask, 0.5 mL of 0.05% trypsin-EDTA (Gibco) were added and the flask was placed back in the incubator for two minutes. To deactivate the trypsin, 2 mL of cell medium was added to the flask and the cells were thoroughly mixed until all visible aggregates were debulked. The resulting cell suspension was directly used for the loading onto the chip. Primary mouse retina pigment epithelium (RPE) were a gift from Sabrina Carrella, PhD (TIGEM) and were obtained as previously described [Gibbs and Williams, 2003].

### Loading of the cells onto the microfluidic chip and static cell culture

To ease the wetting process, the temporary hydrophilicity of the PDMS surface after the plasma activation was exploited. Therefore, the wetting of the channels was carried out within minutes after the bonding procedure. The wetting and the cell loading operations were carried out upon visual check using an inverted stereomicroscope (Leica Microsystems). For the wetting of the microfluidic channels, 10 μL of PBS were added in port B of the chip until all channels were filled. If required, all ports were filled with 10 μL of PBS and the chip was placed in a desiccator, vacuum was applied for 15 minutes or until all air bubbles disappeared. Successively, the PBS was removed from all reservoirs and 10 μL of cell suspension (1–5·10^6^ cell/mL) were added to port B of the device. Cells started to flow through the main channel and to enter the chambers (**Error! Reference source not found.**b). When a given chamber was filled with the suited number of cells, 10 μL of cell medium were added to the respective port to decrease the flow rate across that chamber. For the cell lines used in this work (HeLa, CHO, AU565 and HCC38 cells) it was found that between 2 to 20 cells were sufficient to sustain their proliferation in a chamber. The initial quantity of cells to load into the chambers can vary according to the experimental requirements and the cell line. This parameter can be easily adjusted by varying the cell suspension concentration and the loading time. To avoid adhesion of the cells to regions of the chip other than the chambers (e.g. main channel, reservoirs) it is preferable, however, to adjust the cell concentration while keeping the loading time within a few minutes. When all chambers were filled, 20 μL of cell media was added to port A and port B was emptied to wash the main channel from undesired cells. Port B was then rinsed and filled with 20 μL of cell media. Next, all input ports from #1 to #5 were filled up to a final volume of 20 μL of fresh cell media. An equal volume of cell media in all ports prevents the formation of pressure drops across the chip and enables the static cell culture. Ultimately, 2.5 μL of mineral oil for cell culture (M8410, Merck kGaA) were added to each reservoir to prevent the evaporation of its content when the chip was placed into the cell incubator. The chips stayed in the incubator overnight at 37°C, 100% humidity and 5% of CO_2_ atmosphere prior to the beginning of the experiments.

### Microscopy acquisitions

Live cell widefield microscopy data was acquired via a Nikon Ti Eclipse microscope equipped with a mercury lamp (Intensilight, Nikon), a EMCCD digital camera (iXon Ultra 897, Andor Technology Ltd) and an incubation chamber (H201-OP R2, Okolab). Prior to the mounting of the chip, the incubator was equilibrated to a temperature of 37°C and 5% CO_2_ humidified air. Time-lapse acquisitions up to 20 hours were acquired using a 40x air objective (CFI Plan Fluor DLL 40x, 0.75 NA, Nikon Instruments) to collect the largest amount of signal. However, because of the off-center position of chamber #5 in this specific device, the acquisition of this chamber with this objective was not achievable in our microscope. For the single cell analysis, the fluorescence of the cells from all chambers was acquired at the end of the 20 hour treatment using a 20x air objective (CFI Plan Fluor DLL 20x, 0.5 NA, Nikon Instruments). Depending on the experiment, images were collected in phase contrast (PC) and epifluorescence for green, red and yellow wavelengths using the exposure times and filter sets respectively reported in Table 1.

**Table 1.** Summary of the microscopy settings used in this work.

|  | **Exp time (ms)** | **Ex (nm)** | **Dm (nm)** | **Em (nm)** |
| --- | --- | --- | --- | --- |
| **Phase contrast** | 300 | -- |  | -- |
| **Chroma EGFP** | 300 | 470/40 | 495 | 515/30 |
| **Nikon TRITC** | 100 | 540/25 | 565 | 605/55 |
| **Nikon FITC** | 300 | 480/30 | 505 | 535/45 |

### Multi-input configuration: stress response of CHO-K1 cells under tunicamycin stimulus

After overnight seeding, VersaLive chips with CHO-K1 cells in static culture were removed from the incubator. To each chip, reservoirs A and B were emptied. Next, the content of reservoirs #2 and #4 was renewed with 15 μL of fresh F12 cell medium. Then, the content of reservoirs #1, #3 and #5 was replaced with 15 μL of tunicamycin 0.5 μg/mL in F12 cell medium. Then, 1 μL of sulforhodamine B 0.1 mg/mL (Merck kGaA) was added to reservoir #3 as flow tracer. To all filled reservoirs (#1 to #5), 2.5 μL of mineral oil for cell culture were added to prevent their evaporation. Reservoirs A and B were left empty. The chip was then mounted on the microscope for live cell imaging for a time lapse acquisition of 20 hours at intervals of 15 minutes between each acquisition.

### On-chip immunostaining of AU565 and HCC38 breast cancer cells

AU565 and HCC38 breast cancer cell lines were loaded on VersaLive chips and grown overnight in static cell culture as described in a previous section of this work. For the on-chip chemical fixation, after removal of the cell medium, all ports were washed with 20 μL of PBS. Cells were perfused in multi-input mode with paraformaldehyde (PFA, 4%_w/v_ in PBS) for 10 minutes. Next, all ports were washed twice with 20 μL of PBS followed by 5 minutes of perfusion of PBS in multi-input mode. To quench the fluorescence of PFA, a blocking solution (50 mM NH_4_Cl; 0.5% BSA in PBS) was perfused in multi-input mode for 30 minutes. Ultimately, all ports were washed with 20 μL of PBS. For the on-chip immunostaining, anti-HER2 antibody (BB700 Mouse Anti-Human Her2/Neu, BD OptiBuild) diluted in blocking buffer was perfused in multi-input for a time defined by the experiment (10 or 30 minutes). Then, all ports were washed twice with 20 μL of PBS. To preserve the sample, all ports were filled with 20 μL of PBS and 2.5 μL of mineral oil for cell culture to prevent evaporation. Samples were stored at 4°C. To quantify the stress response of CHO-K1 cells treated and untreated with tunicamycin, live cell microscopy images were analyzed using the Trainable WEKA Segmentation [Arganda-Carreras et al., 2017] plugin in ImageJ software. In brief, the software was trained in distinguishing CHO cells within the culture chambers of VersaLive. The probability map returned by the software was thresholded to be converted in a segmented mask. The mask was then superimposed onto the original image. The average intensities of the resulting single cells were used to quantify their stress response.

### On-chip immunostaining of primary mouse RPE cells

Primary mouse RPE cells were loaded on VersaLive chips and let adhere to the microfluidic chip surface overnight in static cell culture, as described in a previous section of this work. On-chip chemical fixation was carried out as previously described for the cancer cell lines. Immunostaining on chip was initiated by perfusing for 10 minutes the permeabilization buffer (0.3% Triton X-100, 5% FBS in PBS) in multi-input mode. Next, the anti-citrate synthase primary antibody (ab96600, Abcam) in blocking buffer (0.5% BSA, 0.05% saponin, 50 mM NH_4_Cl, 0.02% NaN_3_ in PBS) was perfused in multi-input mode for 10 minutes. All ports were then washed with 20 μL of PBS. Successively, fluorescently labeled donkey anti-rabbit Alexa Fluor 488 (A-21206, Thermo-Fisher Scientific), Alexa Fluor 568 phalloidin (A-12380, Thermo Fisher Scientific) and DAPI (D1306, Thermo Fisher Scientific) were perfused for 10 minutes in multi-input mode to stain mitochondria, f-actin and nuclei, respectively. All ports were then washed twice with 20 μL of PBS. To preserve the sample, all ports were filled with 20 μL of PBS and 2.5 μL of mineral oil for cell culture to prevent evaporation. Samples were stored at 4°C.

### Evaporation of the inlet reservoirs

The effect of the evaporation at the inlet reservoirs was assessed by running two different microfluidic devices in multi-input mode. Input reservoirs #1, #2, #4 and #5 were filled with 25 μL of DI water; input reservoir #3 was filled with 25 μL of rhodamine B (0.1 mg/mL). For one of the two devices, 2.5 μL of mineral oil for cell culture were added to the inlet reservoirs to prevent their evaporation. The intensity of rhodamine B was measured within the culture chamber over 15 hours at 15 minutes intervals. Measurements were performed at the same experimental conditions used for live cell microscopy experiments. Data was processed using ImageJ software.

### Sequential on-chip cell lysis

For the sequential on-chip cell lysis, a VersaLive chip with HeLa wild type cells grown in multi-input mode was used. The composition of the lysis buffer used in this work was adapted from literature [Macosko *et al.*, 2015] to be 200 mM Tris pH 7.5 (Merck kGaA), 6% Ficoll PM-400 (Merck kGaA), 0.2% Sarkosyl (Merck kGaA), 20 mM EDTA (Thermo Fisher Scientific). At each iteration of lysis, the content of the reservoir relative to the target chamber was removed and then replaced with an equal amount of lysis buffer. For every exchange, the chip was removed from the microscope stage and then placed back onto it once the reservoir was filled with lysis buffer. Then, a time lapse of the lysis was recorded in phase contrast at 12 frames per minute.

### Chip removal upon reversible bonding

To facilitate the removal of the PDMS, a single-edge razor blade was gently inserted at the interface between the PDMS and the glass slide all along the perimeter of the chip. Successively, the PDMS chip was pinched to peel it off the glass slide that sealed the channels. For the out-chip chemical fixation, the PDMS chip was first removed from the glass slide. Then, the glass slide with the exposed cells was rinsed with PBS and later immerged in the 4%_wt_ PFA solution for 30 minutes at room temperature. The slide was then rinsed with PBS and gently dried.

---

**Supplementary Figure 1.**
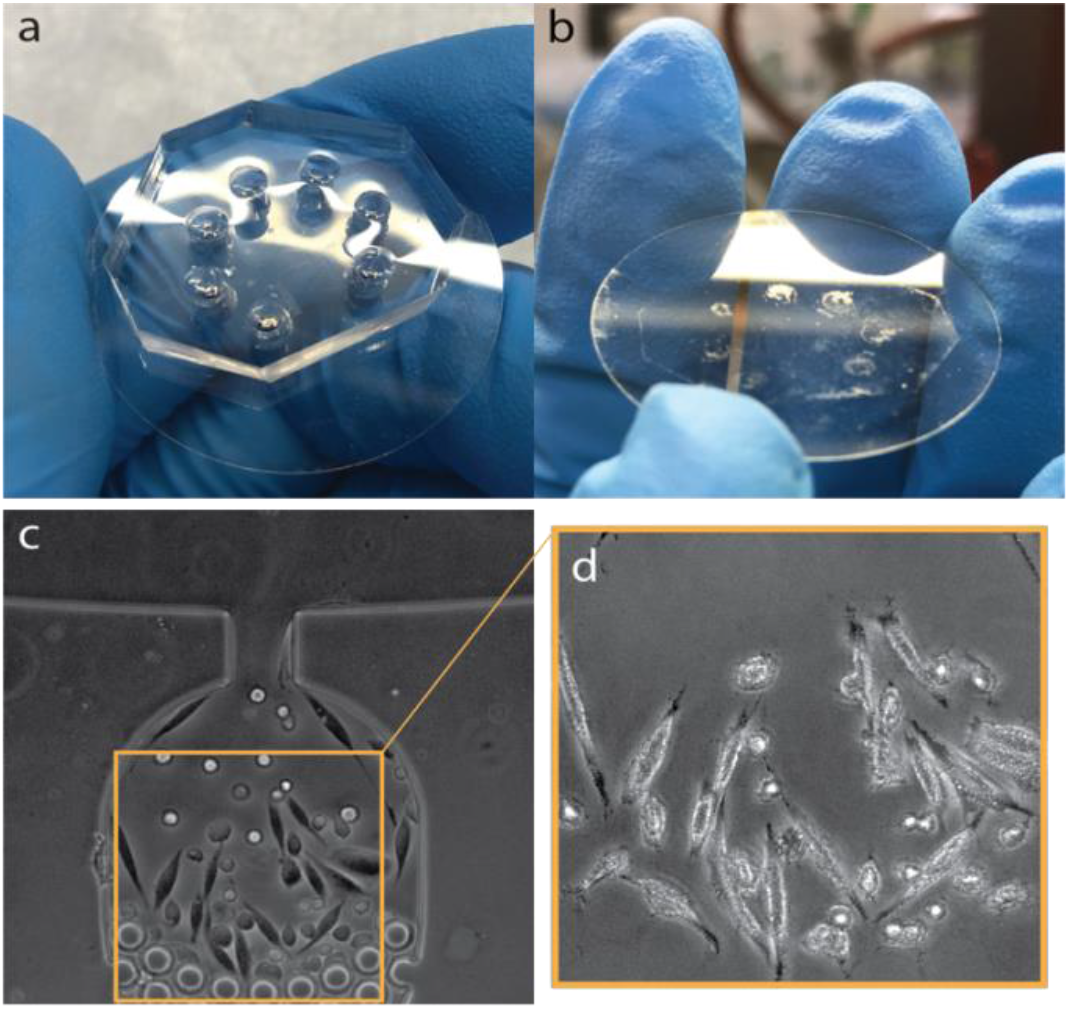
(a, b) Opening of the chip for direct access to the cultivated cells. (c, d) On-chip chemical fixation of previously live-imaged cells and PDMS chip removal. PDMS removal does not affect the cell position of on-chip chemically fixed CHO-K1 cells respect to the live imaged sample. We envision this feature to enable cell re-plating, cell sequencing and all applications that require full recovery of the sample.

**Supplementary Figure 2.**
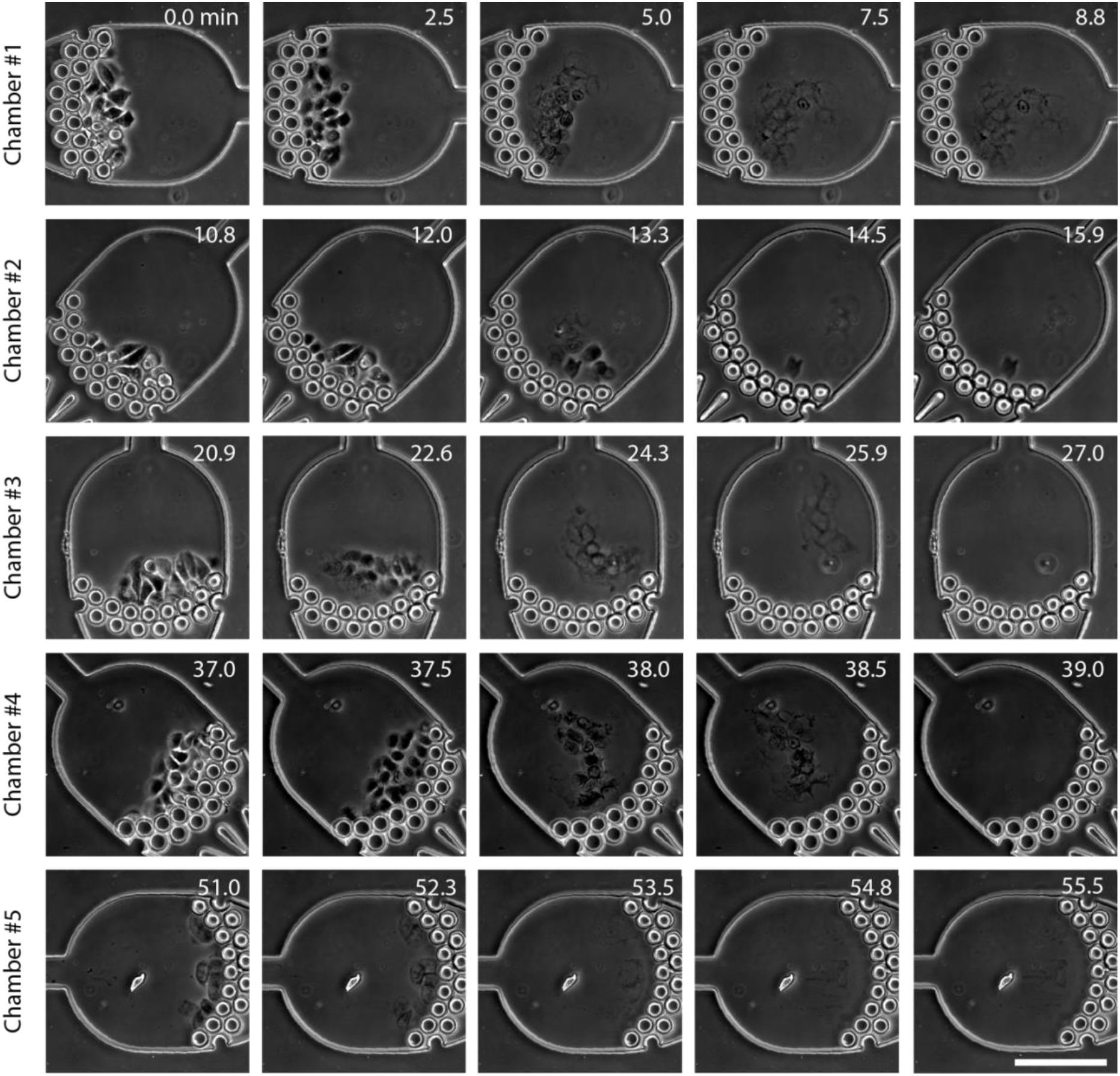
Selective lysis of the chamber content in VersaLive. HeLa cells were grown onto the platform in multi-input mode and later selectively dissolved by exchanging the cell medium with lysis buffer. As appreciable from the frames, cells in the chambers are not perturbed by the process until the beginning of their lysis process. Scale bar 150 μm.

## Acknowledgments

This work was supported in part by the COSY-BIO (Control Engineering of Biological Systems for Reliable Synthetic Biology Applications) project, which has received funding from the European Union’s Horizon 2020 research and innovation program (grant agreement 766840), in part by “OPL-APPS - IIoT OPEN Platform e Applicazioni per il manufacturing” (grant PON ARS01_00615) and in part by BrightFocus Foundation (grant n. M2020184).

## Author Contributions

GN designed the device and performed experiments. GV, SB and SC performed immunostaining experiments. DdB conceived the idea and supervised the work. GN and DdB wrote the mansucript.

